# Homeostatic Forces Shaping the Daily Pattern of Sleep Propensity

**DOI:** 10.1101/2025.03.09.642173

**Authors:** Vasili Kharchenko, Michael Rozman, Arcady A. Putilov, Irina V. Zhdanova

## Abstract

The discrepancy between high sleep need and the ability to initiate sleep underlies insomnia, the most prevalent sleep disorder, whose nature remains obscure. Sleep need increases monotonically with prolonged wakefulness, as reflected in the rising intensity of sleep after varying wake intervals. In contrast, sleep propensity – the ability to transition from wake to sleep – follows a bimodal pattern, peaking in the mid-afternoon, dipping in the evening, and rising again near bedtime. Previously, we demonstrated that sleep structure dynamics can be effectively modeled using probability waves. Here, we extend this wave model of homeostatic regulation of sleep to the period of wakefulness and show that its extrapolation predicts the bimodal pattern of wake-to-sleep transitions. This pattern arises from the interplay of two key factors in state transitions: wake-state instability and interaction strength between states. While wake-state instability increases monotonically, interaction strength follows a bimodal pattern. Their combined effect produces a bimodal probability of state transitions, aligning closely with experimental data. The mid-afternoon peak corresponds to maximal interaction at the homeostatic equilibrium of sleep and wake states, whereas the evening dip reflects minimal interaction, counteracting high wake-state instability. An exponential rise in both factors by the end of the day facilitates sleep onset at bedtime. Our experimental findings on sleep deprivation support the model predictions. Understanding the relationship between sleep need and ability to initiate sleep may offer valuable insights for optimizing daytime performance and sleep quality in both health and disease.

**Statement of significance:** Monotonic increase in sleep need throughout the day contrasts sharply with the bimodal pattern of the ability to fall asleep, referred to as sleep propensity. This phenomenon, inherent to normal sleep physiology, may also underlie major insomnias – inability to fall asleep despite high sleep need. We applied the wave model of sleep dynamics to unveil how the bimodal daily sleep propensity pattern can emerge from the non-linear changes in homeostatic strength of interaction between the wake and sleep states.

## INTRODUCTION

Animals alternate between two primary physiological states: wake and sleep. These states follow predictable, species-specific patterns and are regulated by both homeostatic^1^ and circadian^2^ mechanisms. In humans, the likelihood of transitioning from wake to sleep – referred to as sleep propensity – typically follows a distinct bimodal pattern (Fig. 1a) ^3,4^. Experimentally, sleep propensity is assessed by measuring sleep onset latency after varying durations of prior wakefulness or time spent asleep over a brief sleep opportunity. Such short-sleep protocols show^4^, that, after morning awakening, sleep propensity initially declines for 1–2 hours. It then gradually increases, peaking slightly in the mid-afternoon, before decreasing to a daily minimum in the early evening. Shortly before habitual bedtime, it rises again.

**Figure 1.**
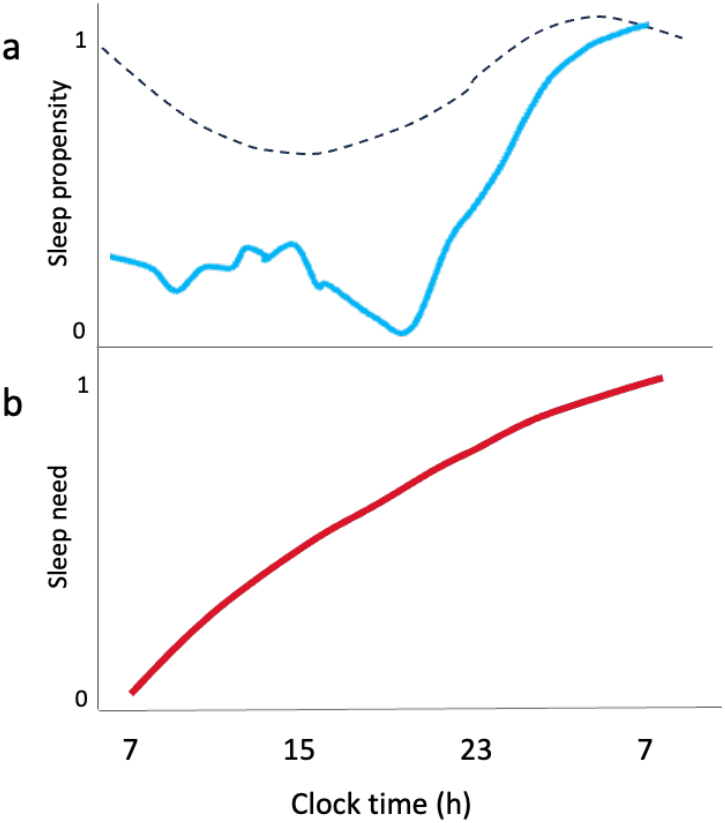
Difference in daily patterns of sleep need and sleep propensity in humans. **a** Schematics of the bimodal pattern of sleep propensity in the absence of nighttime sleep, as based on ultra-short sleep-wake cycles data^4^. Circadian modulation of sleep state (dashed line), based on the inverted circadian pattern of core body temperature^2^. The clock suppresses sleep during the day and facilitates it at night. **b** Schematics of exponential rise in sleep need, based on the intensity of the initial sleep episode^1^.

This pattern differs sharply from the continuous rise in sleep need^1^ during wakefulness. It is quantified based on the initial intensity of sleep or based on the cognitive decline measures that indicate wake state instability^5^. Sleep need increases monotonically over the day and follows a negative exponential function^1^ (Fig. 1b).

This gap between the need for sleep and the ability to initiate it has long puzzled researchers. Several hypotheses have been proposed to explain the bimodal pattern of sleep propensity, most of which emphasized the modulatory effects of the circadian clock^6-14^. In humans and other diurnal animals, daytime activity of the circadian clock suppresses wake-to-sleep transitions and promotes wakefulness^14,15^, counteracting the sleep-promoting effects of the rising sleep need (Fig. 1a). This effect declines in the evening, with maximal sleep-promoting effect of the clock manifesting near the end of the night. However, the daily patterns of the clock’s activity and sleep propensity differ notably: the mid-afternoon peak in sleep propensity coincides with the clock’s maximal wake-promoting effect, while the evening dip in sleep propensity appears as the clock’s wake-promoting influence begins to wane (Fig. 1a,b).

Under regular conditions, both homeostatic and circadian influences shape sleep. However, in this study, we use the wave model of sleep dynamics^16^ to focus specifically on the homeostatic regulation of the sleep-wake cycle. We hypothesize that the bimodal daily pattern of transitions between wake and sleep states arises from a nonlinear homeostatic relationship between these states and that two distinct factors govern their interaction. Our approach draws on a well-established principle from physics: state transitions depend on two key factors – the relative instability of one state compared to the other and the strength of interaction between them. Without sufficient interaction, even a highly unstable, high-energy wake state may remain metastable, preventing a transition to a low-energy sleep state. This state interaction, whether direct or indirect, can also be modulated by external factors such as the circadian clock.

The wave model describes sleep-wake cycle as a dynamic homeostatic process governed by two interacting probability waves representing wake and sleep states, each evolving within its Morse potential well^16^. The Morse potentials have been widely used in physics and biophysics to describe state transitions, particularly in systems where energy landscapes involve metastable states and nonlinear dynamics. Fitted to regular sleep patterns, the wave model accurately captures the experimentally observed durations and intensities of consecutive episodes of non-rapid-eye-movement sleep (NREMS) and REMS. Moreover, the model correctly predicted an invariant relationship between NREM and REM sleep, which was experimentally validated, along with the effects of acute sleep deprivation and excess sleep on subsequent sleep structure^16^.

Here, we demonstrate that the bimodal pattern of the probability of state transitions predicted by the wave model – considering only the homeostatic relationship between wake and sleep – closely aligns with the experimentally observed bimodal sleep propensity curve. This alignment emerges from bimodal variations in the strength of interaction between wake and sleep states. Moreover, we demonstrate that this pattern is altered by sleep deficits of varying severity, fully aligning with the model’s predictions.

## RESULTS

### Transitions between sleep and wake states in the wave model of sleep

The wave model’s fit on the duration of consecutive NREMS and REMS episodes in normal sleep^16^ allows it to track in parallel the dynamics of the two critical factors governing state transitions: the instability of the wake state relative to the sleep state, and the strength of interaction between these states. Figure 2a illustrates how the model represents wake and sleep states as two interacting Morse potential curves (*U*_*W*_ and *U*_*S*_) and describes their behavior as two respective probability waves.

**Figure 2.**
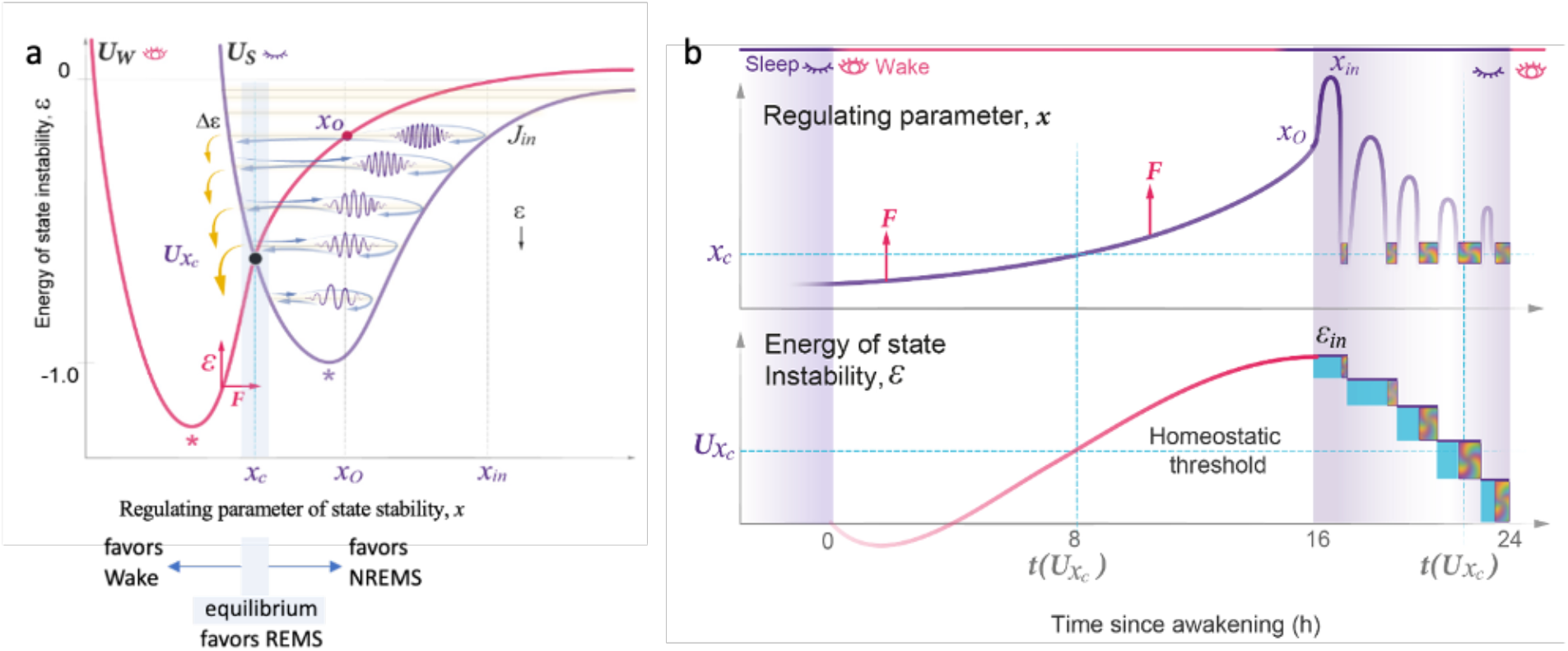
The schematics of the wave model of sleep dynamics. a. In the wake state (*U*_*W*_, red potential), the driving force *F* (horizontal red arrow) of homeostatic, circadian or environmental nature increases the value of the regulating parameter *x* (horizontal axis) beyond the sleep-wake homeostatic equilibrium setpoint *x*_*c*_ (vertical cyan line) and the *x*_*c*_ region of efficient interaction of two states (gray area). This rises the energy of state instability *ε* (vertical red arrow and vertical axis) above the homeostatic energy threshold (*Ux*_*c*_, black dot; the crossing point of the two potential curves). Sleep (*U*_*S*_, purple potential) is initiated at *x*_*o*_ (red dot) of the initial energy level *J*_*in*_ (first sleep cycle). Relaxation of *x* and *ε* occurs in the form of sleep wavepacket propagating along energy levels (blue arrows), where *x*_*in*_ – maximal deviation achieved during the first sleep cycle. At the end of each cycle, around *x*_*c*_, a portion of the wavepacket energy Δ*ε* (yellow arrows) is released. Stars– maximal stability of the corresponding state. The schematics serves to only illustrate the concept. For actual position of *Ux*_*c*_ and *J*_*in*_ within *U*_*S*_ see Fig. 3a. The figure is taken from^16^, with permission. **b. *Top panel:*** Time-dependent changes in the regulating parameter of state stability *x* (vertical axis) over wake (white area) and sleep (purple area). In sleep, the period of *x* oscillations corresponds to the duration of consecutive NREMS episodes, while a decline in their amplitude squared corresponds to the drop in NREMS intensity. Each cycle ends around *x*_*c*_ (horizontal cyan line), the region of sleep–wake equilibrium where REMS episodes occur (multicolor blocks). *F*—the driving force (red arrow) that increases *x*. ***Bottom panel:*** Time-dependent increase in the energy of state instability *ε* (vertical axis) over the wake state (white area) and stepwise decline over consecutive sleep cycles (block width—duration; NREMS—cyan; REMS—multicolor). Small reduction in *ε* at the start of wake period corresponds to sleep inertia. Maximal energy level reached 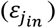 defines initial NREMS intensity. Linearly increasing portions of energy released Δ*ε* correspond to REMS intensity (block height). Horizontal cyan line—potential energy of homeostatic threshold, *U(x*_*c*_). Vertical cyan line—timing of *U(x*_*c*_), with daytime timing around mid-afternoon peak in sleep propensity. Horizontal axis—time since awakening (hours). The figure is taken from^16^, with permission.

The sleep-wake cycle is a homeostatic process in which deviations from equilibrium activate compensatory mechanisms when one of the alternative states becomes highly unstable. The model revolves around a single variable, the regulating parameter of state stability (*x*), which serves as a one-dimensional representation of the overall physiological state and indicates the system’s capacity to maintain stability. This parameter responds to various influences, including homeostatic, circadian, and environmental forces. Its fluctuations define the energy of state instability (ε), which rises during wakefulness and declines during sleep (Fig. 2b).

At the point of intersection of the two potentials (*Ux*_*c*_), the two states achieve homeostatic equilibrium where the energies of the wake and sleep states are equal. Thus, the *x*_*c*_ region marks the homeostatic equilibrium threshold in state instability beyond which one state becomes more stable than the other. The lower the energy of the state, when compared to another, the relatively more stable it is, as per the minimal energy principle. Thus, at *x* < *x*_*c*_, the wake state exhibits greater stability, whereas at *x* > *x*_*c*_, the sleep state becomes more stable (Fig. 2a,b).

The normal sleep architecture based on the data collected in young healthy individuals was defined by the simple characteristics of the *U*_*S*_ potential well – its width and the initial energy level at which sleep is initiated (*J*_*in*_) – and the energy of homeostatic equilibrium (*Ux*_*c*_) where the *U*_*S*_ and *U*_*W*_ potentials cross (Fig. 2a,b). These were estimated through the model analysis of just two experimental variables – the duration of consecutive NREMS and REMS episodes, respectively^16^. Once known, these characteristics allowed us to determine the two model parameters on which the probability of state transition relies: state instability (*ε*) and the strength of interaction (*Z)* between the sleep and wake states, as detailed in the Methods section.

Figure 2b illustrates that state instability, represented by ε and *x*, increases during the wake state. This is except for a brief period when ε continues to decline immediately following awakening, a consequence of the typical inertia of the damping forces (i.e., the short-lived carryover of low instability from the prior sleep state). Figure 3 illustrates the *x*-dependent dynamics of ε in parallel with the second parameter of interest, the strength of interaction between the two states *Z*. When the *U*_*S*_ and *U*_*W*_ potential curves are close to one another, the interaction between the two states becomes stronger, leading to an increased likelihood of state transitions. Notably, the potential curves are maximally close to each other near the homeostatic equilibrium point (*Ux*_*c*_). Below *Ux*_*c*_, the proximity between the curves decreases, whereas above *Ux*_*c*_, the proximity of the two states demonstrates a non-linear relationship with *x*. Specifically, the proximity between the two states is first declining and then increasing again over the period of wakefulness (Fig. 3a). This gives rise to a bimodal *Z* pattern (Fig. 3b).

**Figure 3.**
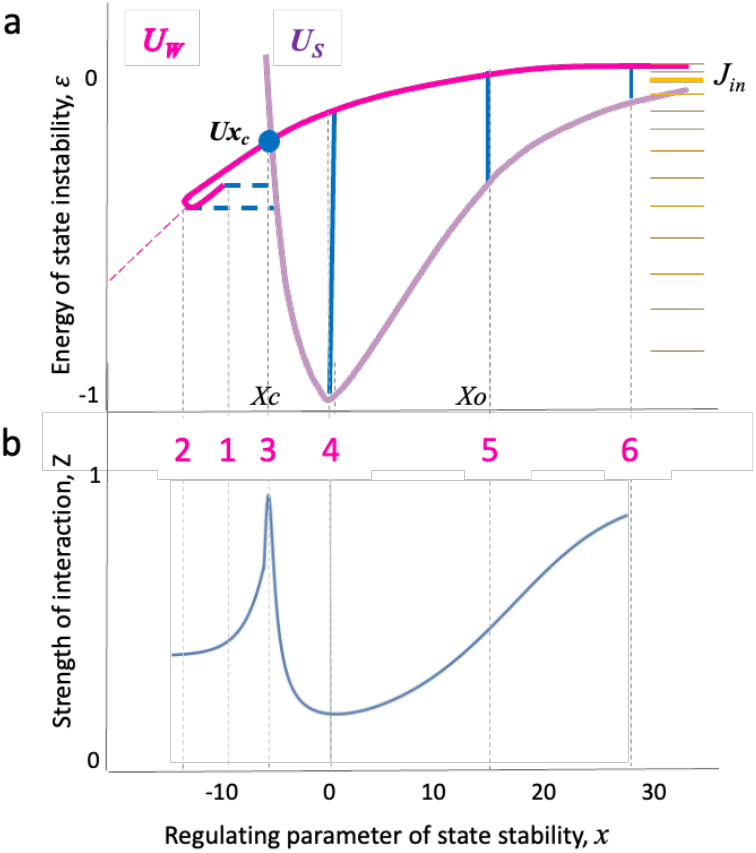
Temporal relationship between the wake state instability and strength of interaction between the sleep and wake states. **a** The dynamics of the energy of state instability, *ε* (magenta line) depends on the regulating parameter of state stability *x* (horizontal axis). Sleep-propensity zones: the numbers (1-6 in pink) and corresponding thin vertical lines indicate points of comparison between *ε* and the proximity (blue lines) between sleep (*U*_*S*_, purple) and wake (*U*_*W*_, magenta) potential curves (as in Fig. 2). Following morning awakening (1), the energy of state instability continues to decline (2, sleep inertia), then rises towards the homeostatic equilibrium (3, *X*_*c*_ and *Ux*_*c*_, blue dot) and beyond it (4-6) during consolidated wakefulness. The point *X*_*o*_ (5) - habitual sleep onset at the initial energy level *J*_*in*_, corresponding to the first sleep cycle. Point 6 –increase in state instability and the proximity of two states during sleep deprivation. Yellow lines illustrate gradual increase in energy gaps of the Morse potential towards low energy levels. As detailed in^16^, on average, regular sleep starts around level 10 and *Ux*_*c*_ is at around level 7 in young adults with normal sleep. **b** The strength of interaction (*Z*, blue line) between sleep and wake states positively correlates with *x* and proximity between the two states, thus negatively correlates with the distance between the potential curves (blue lines in Fig. 3a). Following morning awakening (point 1), *Z* declines (point 2, sleep inertia) and then increases towards the crossing of the two potential curves *Xc*, where the proximity is maximal, while the two states are at equilibrium (point 3). Further rise in state instability is associated with large distance or minimal proximity between the two states (point 4), hence, with weak strength of their interaction, which then gradually increases towards the point of habitual sleep onset *Xo* (point 5) and further (point 6). If sleep is prevented (point 6, after the whole-night sleep deprivation), *Z* continues to rise due to the rise in state instability and growing proximity between the potential curves at high *x*.

The mathematical analysis of these two parameters, ε and *Z*, follows a continuous function (see Methods). In Figure 3, we illustrate their dynamics using six key points, designated as sleep-propensity zones for further analysis of the probability of state transitions. Point 1 represents awakening or the onset of the daily wake period, followed by a relaxation of state instability due to the inertia of the damping forces (Fig. 3a). This minor decrease in state instability is associated with a reduction in the proximity of the two states (point 2). Subsequently, the driving forces of homeostatic, circadian, and environmental nature favor the wake state and elevate state instability towards the homeostatic equilibrium threshold (*x*_*c*_, point 3). At this threshold, the potential energies of the two states are equalized (*Ux*_*c*_), their proximity reaches its maximum and Z reaches its peak (Fig. 3b).

Beyond the *x*_*c*_ equilibrium region, the rise in state instability favors the transition from the increasingly unstable wake state to the more stable sleep state (Fig. 3a). However, this is accompanied by a significant reduction in the proximity of the potential curves representing these two states, reaching its maximum distance and thus minimal strength of interaction *Z* at point 4 (Fig. 3b), which makes transitioning from wake to sleep difficult. Once point 4 is surpassed, the further increase in the energy of state instability ϵ brings the two potential curves closer together (points 5 and 6), thus increasing *Z*. Consequently, both critical parameters – state instability and strength of interaction - now favor the transition from wake to sleep.

### Wave model predicts the bimodal pattern of sleep propensity

Using the wave model fit on the experimental observations of NREM and REM episode durations in young, healthy males, we then examined its predictions of how the probability of transitioning into sleep state depends on the time since awakening. The 24-hour dynamics of *x* and *ε*, as predicted by the model (Fig. 2b), along with the established correspondence between *x, ε* and *Z* (Fig. 3), enabled us to reconstruct the daily pattern for the probability of state transition (*P*) (see Methods). The overall bimodal profile and the timing of its specific peaks and dips in the absence of sleep (Fig. 4a) closely resembled the profile of sleep propensity documented experimentally using an ultra-short sleep protocol^4^ (Fig. 1a).

**Figure 4.**
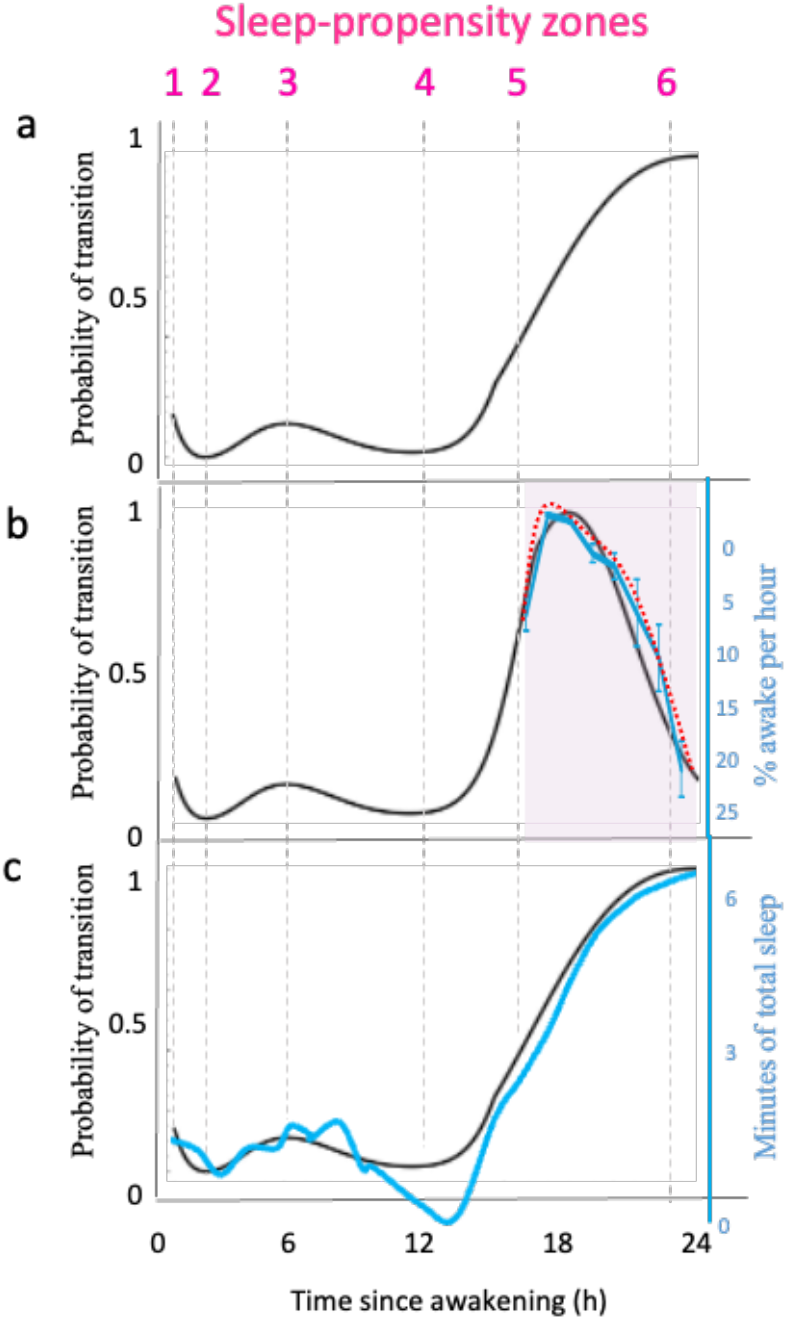
Daily variation in the homeostatic pattern of state instability and probability of sleep-wake state transitions. a. Model curve for the homeostatic component of the probability of state transition (sleep propensity) in the absence of overnight sleep. Sleep-propensity zones (pink numbers) as in Fig. 3. b. Homeostatic component of sleep propensity during the day (white area) and over an 8-h nighttime sleep (purple area). Black – model prediction based on consecutive 0.1-hour intervals; cyan curve – experimental data for mean % time (SEM) spent awake per hour of overnight sleep in young healthy males (n=39 nights); red dashed line – sleep-dependent variation in % time spent awake (inverted curve), schematic based on Dijk & Czeisler^7^. Horizontal axis – time since awakening (hours). c. Comparison of model (black) and experimental (cyan) curves of probability of state transitions. In the absence of sleep (model) or during short-sleep protocol (13 min-wake/7-min sleep opportunity), schematics based on ^4^.

Specifically, the morning dip in sleep propensity corresponded to minimal wake state instability following the period of sleep inertia (point 2). On the other hand, the mid-afternoon ‘siesta’ peak aligned with the homeostatic equilibrium and maximal proximity of the two states at *Ux*_*c*_ (point 3). The evening dip in sleep propensity was within the region of low interaction between the two waves (point 4). The surge in sleep propensity at the habitual sleep onset time, known as the ‘sleep gate’^4^, corresponded to the rapid increase in *P* at point 5 due to the surge in both the states’ interaction and energy of state instability. In the absence of overnight sleep (Fig. 4a), further rise in the probability *P* of the wake-to-sleep transition corresponded to the continued exponential increase in ε and *Z* around point 6 (Fig. 3a).

Figure 4b demonstrates the result of the wave model’s approach to estimating sleep propensity during sleep. It predicts that sleep propensity changes with each sleep cycle, specifically declining after each REM episode, which is associated with a reduction in *ε* (Fig. 2, 3). This decline in sleep propensity depends on the ratio of consecutive NREM and REM sleep episode durations, as described in detail in the Methods section.

Experimentally, sleep propensity during sleep is assessed by the percentage of time spent awake after sleep onset — essentially a measure of ‘wake propensity’^7^. To illustrate overnight sleep propensity, Figure 4b presents an inverted wake propensity curve from our study of young males with normal sleep patterns. Both experimental and theoretical approaches yield similar results (Fig. 4b), aligning with earlier findings on sleep-dependent wake propensity changes in forced-desynchrony studies7. The probability of state transitions shows an initial rise in sleep propensity after sleep onset (a period akin to ‘wake inertia’), peaking by the end of the first sleep cycle before monotonically declining thereafter (Fig. 4b).

Documenting sleep propensity requires providing subjects with repeated sleep opportunities, during which they spend some time asleep4. The model predicted that daytime sleep would delay the onset of the mid-afternoon peak and evening dip in sleep propensity, as the energy of instability (or sleep need) would not accumulate during sleep periods. Accordingly, the model’s peaks and dips occurred slightly earlier than those observed experimentally (Fig. 4c). Notably, the evening dip was less pronounced in the model curve, as it accounted only for homeostatic effects, whereas the experimental curve reflected both homeostatic and circadian influences.

### Effects of sleep deficit on the sleep propensity profile

Previously, the wave model accurately predicted the effects of acute sleep deprivation on sleep architecture by accounting for the increase in wake state instability^16^. Sleep initiation at a higher energy level of the *U*_*S*_ curve (Fig. 2a) explained not only the characteristic higher intensity of NREM sleep but also the reduction in both the duration and intensity of the initial REM sleep episodes following sleep deprivation. By incorporating an additional parameter - the strength of interaction between the two states (Fig. 3b) - we now addressed the changes in sleep propensity following sleep deprivation.

We analyzed the effects of sleep deprivation resulting in varying degrees of sleep deficit and residual wake state instability, as reflected by greater deviations in ε and *x* at the onset of sleep propensity assessment (Fig. 5a). The model predicts that more severe sleep deficits produce more pronounced changes in the daily sleep propensity profile. Mild sleep deficits result in wakefulness initiated near the homeostatic equilibrium threshold, *Ux*_*c*_ (points 1–3) (Fig. 5a). This masks the typical mid-afternoon peak, which instead manifests as a modest initial increase in sleep propensity due to higher *ε* and Z. However, the evening dip in sleep propensity remains evident, as the gradual rise in ε moves through the region around *x*=0, characterized by minimal interaction between sleep and wake states (point 4).

**Figure 5.**
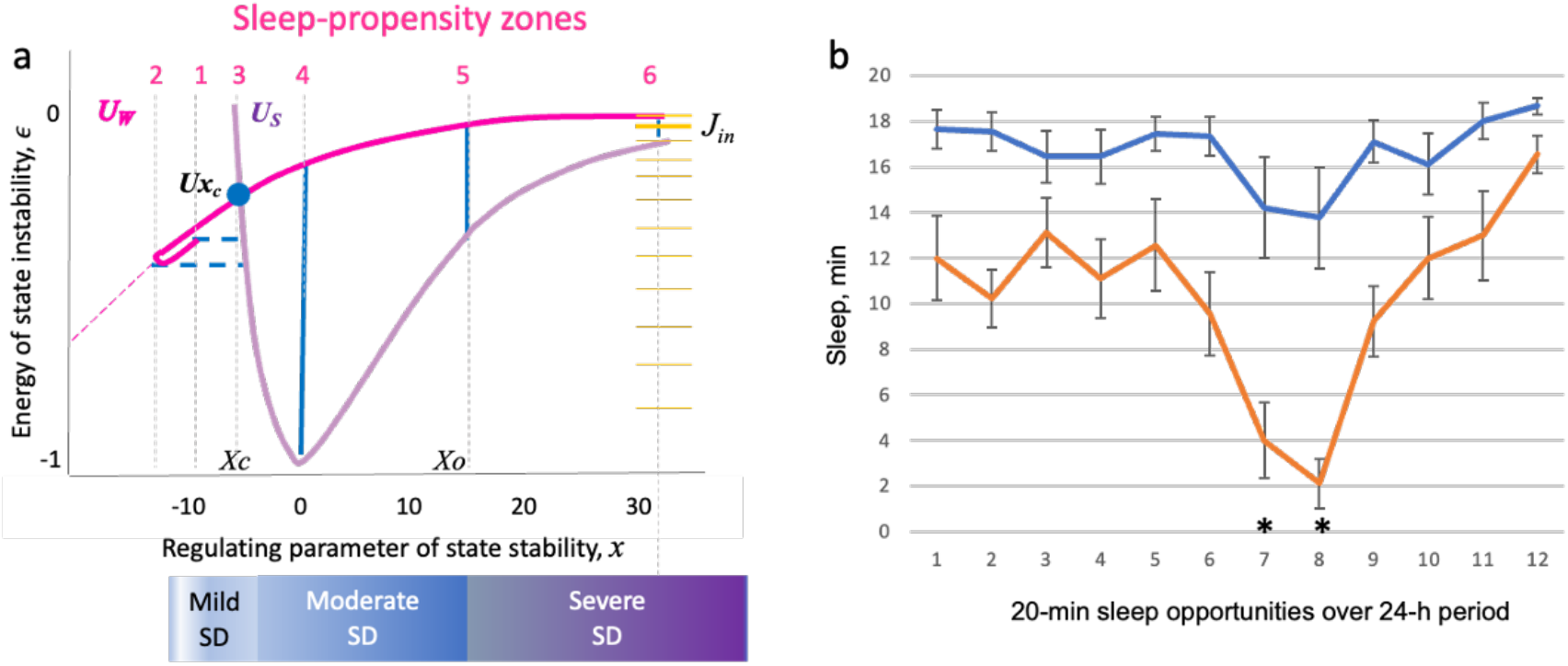
Modifications of sleep propensity profile due to different degree of sleep deficit. a. Under normal sleep conditions, relaxation of the energy of state instability ε during sleep reaches point 2 (pink number). Mild sleep deficit (SD) results in lesser relaxation of ε (points 1-3), masking the mid-afternoon peak, though preserving the evening dip in sleep propensity around point 4. Moderate SD negates the mid-afternoon peak but can also modify the evening dip, since involves the points 3-5 interval of weak interaction between the sleep and wake states. Severe SD negates the entire bimodal pattern of sleep propensity due to a combination of relatively high ε and strength of interaction between states beyond point 5. Sleep-propensity zone 1-6, as in Fig. 3. b. Sleep propensity (minutes asleep during 20-min sleep opportunities provided at 100-min intervals) over 24-h period in subjects with mild SD (red) and severe SD (blue), n=9 per group, * p<0.01.

The model further predicts that a moderate sleep deficit (i.e., sleep propensity measurements begin at *x* values near point 4 of weak interaction between the two states; Fig. 5a), will result in reduced initial sleep propensity despite elevated ε due to low initial *Z*. Subsequently, sleep propensity will continue to rise as it typically does after the evening dip, since both ε and *Z* gradually increase.

In contrast, severe sleep deficits - such as those resulting from overnight sleep deprivation - sustain *x* at levels beyond those associated with typical evening sleep onset (point 5). In this case, sleep propensity assessment begins around point 6, where both state interaction and ε are elevated (Fig. 5a). This combination significantly amplifies sleep propensity and suppresses its normal biphasic variations, as the entire evaluation process takes place within the *x* interval characterized by concurrent increases in state instability ε, regulating parameter *x* and the strength of interaction Z.

To test these predictions, we analyzed sleep propensity data from healthy young male cadets who adhered to a military schedule that provided regular 7-hour nighttime sleep opportunities with no daytime naps^17-19^. Given a recommended sleep duration of around 8 hours per night for young males^20^, this schedule represented a condition of chronic mild sleep restriction. Two groups were studied during a 24-hour short-sleep protocol, consisting of twelve 120-min cycles of alternation of 100-minute wake period with 20-minute polysomnographically monitored sleep opportunity. Prior to the experiment, one group maintained regular 7-hour nighttime sleep (mild sleep deficit), while the other group experienced 24 hours of total sleep deprivation (severe sleep deficit).

Figure 5b illustrates that the daily sleep propensity profiles for both groups aligned with the wave model’s predictions. Consistent with the wave model, the mild sleep deficit group exhibited no discernible mid-afternoon peak, while their evening dip remained prominent. In contrast, the severe sleep deficit group displayed high sleep propensity with no significant variations during the mid-afternoon or evening periods, as was predicted by the model.

## DISCUSSION

Precise mathematical description of a physiological process is more than just a theoretical construct – it serves as a framework that reveals the underlying principles governing that process. It has long been recognized that sleep need does not always correspond to the ease of falling asleep. In his 1992 review^21^, Peretz Lavie, whose lab played a pivotal role in characterizing the daily bimodal profile of sleep propensity^4,22-26^, addressed this issue and noted that a comprehensive model of sleep/wake regulation must integrate at least three aspects of sleep behavior: the speed of sleep onset, sleep duration, and sleep composition. The wave model of sleep dynamics offers such a framework by linking the durations of consecutive NREM and REM sleep episodes with the probability of state transitions, embodying the concept of dynamic homeostasis.

Our analysis shows that wake-state instability and the strength of wake–sleep interaction can be quantified by analyzing changes in overnight NREM and REM sleep episode durations. Previously^16^, we found that fitting the wave model to these experimental parameters – readily available from any polysomnographic record – enables a mathematically accurate representation of normal sleep architecture and yields experimentally validated predictions regarding its modification in response to acute sleep deprivation or sleep abundance^16^. Moreover, the model established an invariant relationship between NREM and REM sleep. This study now demonstrates that such experimental sleep data are also sufficient to uncover the homeostatic component of daily sleep propensity profile. This is analogous to predicting the ease of ascending a hill based on the experience of descending it.

Experimental studies have demonstrated how sleep need^1^ (reflected in the intensity of the initial sleep episode) or wake-state instability^5^ (evident in cognitive deficits) accumulates over the wake period, necessitating a transition from wake to sleep. However, while increased instability of one state relative to another is a necessary condition for transition, it is not sufficient on its own. Low interaction strength or a high activation energy barrier can result in a metastable state, reducing the probability of transition. Using our wave model, we demonstrate how a nonlinear change in the strength of interaction between wake and sleep states can occur and how it can account for the bimodal sleep propensity pattern – even while sleep need increases steadily. The model’s predictions align closely with experimental observations on sleep propensity under habitual sleep-wake conditions, overnight wakefulness, and following both mild and severe sleep restriction.

Our results suggest that, during wakefulness, while state instability increases according to an inverse exponential function, the strength of interaction follows a more complex, bimodal pattern (Fig. 3). It peaks at the homeostatic equilibrium where the two states are in maximal proximity (*Ux*_*c*_), corresponding to the mid-afternoon peak in sleep propensity. The interaction then declines sharply near the minimal energy on the sleep potential curve (*U*_*S*_), which coincides with the evening dip. Later, as the proximity of the two potential curves increases again, interaction strength rises, leading to a surge in sleep propensity during extended wakefulness. Under prolonged sleep deprivation, when state instability is very high, the interaction strength also remains consistently elevated, and the bimodal pattern of sleep propensity disappears.

During each sleep cycle, the strength of interaction fluctuates, being maximal in the *x*_*c*_ region during REM sleep, defining its duration^16^, and lower during NREM sleep. This may explain the higher probability of state transition (awakening) in REM sleep. Over the entire sleep period, state instability decreases with each sleep cycle due to the drop in the energy of state instability. This results in a gradual reduction in sleep propensity, the magnitude of which can be accurately predicted by the model based on the ratio of REM/NREM episode durations. The prediction is consistent with our experimental results and the earlier findings on the percent awake during sleep based on the forced-desynchrony studies^7^.

Beyond the homeostatic process, additional factors – such as circadian, environmental, or pharmacological influences – can also affect the dynamics of sleep propensity. It is therefore essential to determine which factors predominantly influence state instability and which primarily modulate the strength of interaction. The model provides clear predictions that can distinguish these effects based on which specific experimental parameters change.

For instance, the circadian clock plays a pivotal role in sleep regulation. The wave model predicts that if the clock’s effect targets state instability, then the principal measures of sleep architecture (the intensities and durations of NREM and REM episodes, plus sleep onset latency) would vary with circadian phase. In contrast, if the clock modulates the strength of interaction between sleep and wake states, it will primarily affect two measures: REM sleep episode duration and state transitions (i.e., sleep onset latency or sleep propensity). This is because, according to the model, these two sleep measures, but not the other three, depend on the strength of interaction between states^16^. Experimental observations, particularly those in which homeostatic and circadian influences are desynchronized, support this latter scenario, with the clock primarily affecting REM episode duration and sleep propensity^2,6,14,15^. Furthermore, a synergistic homeostatic and circadian effect that reduces interaction strength in the early evening is consistent with the more pronounced evening dip in sleep propensity observed under regular entrained conditions, compared to model-generated curves based solely on homeostatic regulation (Fig. 4c).

We therefore propose that the circadian clock exerts its daytime effect by reducing the strength of interaction between sleep and wake states without directly affecting state instability. Toward the end of the day, as clock activity diminishes, disinhibition of state interactions facilitates a surge in sleep propensity and, along with the homeostatic effect on interaction strength, ultimately promotes sleep onset. This is not to be confused with the indirect effect of the clock, which is by attenuating state transitions during the day prolongs and consolidates wakefulness, thereby indirectly increasing overall wake-state instability by the end of the day.

These analytical capabilities of the wave model can be extended to evaluate how specific pharmacological agents or other interventions (e.g., light, exercise, meals) differentially affect state instability and interaction strength. This is particularly relevant for insomnias that manifest as an impaired ability to initiate or maintain sleep, despite objective signs of increased sleep deficit and wake-state instability^27^.

In conclusion, mathematical modeling is particularly powerful in fields with abundant experimental data yet unclear regulatory logic. Sleep research has amassed extensive molecular and physiological findings – from neuronal activity patterns to genetic influences – but integrating them into a coherent framework remains challenging. Our model helps organize these findings by distinguishing factors that destabilize wake state (state instability) from those that modulate state transitions. This distinction enables systematic mapping of molecular and physiological changes onto sleep dynamics. By providing a precise mathematical depiction, our model can enhance understanding of sleep regulation, inform targeted treatments for sleep disturbances, and improve overall sleep health.

## METHODS

### Documenting normal sleep architecture

Modeling of the normal daily sleep propensity profile was based on the analysis of sleep dynamics in a group of young, healthy volunteers, as described previously^16^. Representative group data for regular sleep were obtained as part of our larger study on the circadian regulation of sleep and hormonal functions (“Multimodal Circadian Rhythm Evaluation”; PI: IVZ, funded by Pfizer Inc.). The study was conducted in accordance with the Declaration of Helsinki on Ethical Principles for Medical Research Involving Human Subjects and was approved by the Boston University Institutional Review Board. All participants provided written informed consent.

Twenty-four young, healthy male volunteers (Mean ± SEM: 24.5 ± 4.4 years; range: 19–34 years) were selected based on self-reported criteria: 7–9 hours of habitual nighttime sleep, minimal (<1.5 h) changes in sleep duration on weekends, no sleep complaints, no history of chronic disorders or regular medications, no recent trans-meridian travel, no drug use, no smoking, and habitual coffee consumption not exceeding 3 cups per day.

Over the two weeks preceding the inpatient phase, sleep–wake cycles were documented using activity monitors (Philips Inc.) and sleep logs. Subjects then spent 3 consecutive nights in the General Clinical Research Center at Boston University School of Medicine. Bedtimes were scheduled individually based on habitual sleep patterns, and subjects were allowed 9 consecutive hours in bed. Sleep was recorded using polysomnography (Nihon Kohden PSG system) following standard techniques, with sleep stages visually scored in consecutive 30-second epochs. To be included in the regular sleep dataset, each sleep night had to have a sleep efficiency of at least 85% and show no evidence of sleep apnea or other sleep disorders (n = 39 nights total).

NREMS–REMS cycles were defined by the succession of an NREMS episode (minimum duration: 10 minutes) followed by a REMS episode (minimum duration: 3 minutes). No minimum criterion for REMS duration was applied for the final cycle. An NREMS episode was defined as the interval between the first two epochs of stage 2 and the first occurrence of REMS within a cycle, whereas a REMS episode was defined as the interval between two consecutive NREMS episodes or as the interval between the last NREMS episode and final awakening. Intermittent awakenings were not included in the NREMS or REMS periods and were instead quantified as a percentage of wakefulness per sleep cycle or per time unit.

### Documenting Sleep Propensity Following Mild and Severe Sleep Deficits

Sleep propensity was analyzed in 18 male cadets (age: 18–22 years) from the Novosibirsk military school. The experiments conformed to the ethical standards of the Declaration of Helsinki and were approved by the Ethics Committee of the Siberian Branch of the Russian Academy of Medical Sciences. Informed written consent was obtained from all participants.

Prior to the experimental period, cadets followed a military schedule providing regular 7-hour nighttime sleep opportunities with no daytime naps. Given that the recommended average sleep duration for young males is 8 hours^26^, this schedule represented a state of chronic mild sleep restriction (mild sleep deficit). Group 1 (mild sleep deficit, n = 9) maintained their regular sleep schedule, while Group 2 (severe sleep deficit, n = 9) was kept awake for the entire night preceding the experimental procedures. Both groups were studied in parallel.

The detailed experimental protocol and results for sleep measures other than sleep propensity were reported previously^23–25^. Briefly, during a 24-hour experimental period starting at 06:00 h, subjects underwent testing under sleep laboratory conditions using a short-sleep paradigm consisting of 20-minute sleep opportunities interspersed with 100-minute wake periods. Polysomnographic data were collected using a Medicor polygraph (EEG8S, Micromed, Hungary), and sleep stages were scored in 30-second epochs. Sleep initiation was defined as the occurrence of two consecutive epochs of stage 2. Sleep propensity was quantified based on either the total sleep duration during each 20-minute sleep opportunity or the latency to stage 2 sleep.

### Mathematical Modeling

In accordance with the wave model of sleep dynamics^16^, the daily sleep–wake cycle is viewed as the interplay between two interacting probability waves representing the Sleep (S) and Wake (W) states. This model accurately describes four key polysomnographic (PSG) measures, including the durations and intensities of consecutive NREMS and REMS episodes^16^. Through quantitative analysis of overnight sleep architecture, the model identifies parameters influencing state transitions—namely, state instability and the strength of interaction between the two states—which together determine the probability of transitions between S and W states.

Both S and W states arise from the coherent interplay of numerous underlying homeostatic processes. Within the wave model, these underlying processes are represented by S and W wavefunctions, *ψ*_*S*_(*ξ, x*) and *ψ*_*W*_(*ξ, x*) respectively, where ξ denotes the essential set of internal parameters characterizing these processes. The wave functions are normalized to unity, i.e. < *ψ*_*i*_(*ξ, x*)|*ψ*_*i*_(*ξ, x*) > = 1 for i=s,w. The orthogonality < *ψ*_*S*_(*ξ, x*)|*ψ*_*W*_(*ξ, x*) >= 0 of the S and W wavefunctions emphasizes their fundamental internal differences. The functions *ψ*_*i*_(*ξ, x*) can be represented as eigenfunctions that include only real parts.

In our modeling, the single regulating parameter of state stability *x* is employed, and its value defines the specific forms of the two wavefunctions. This parameter is a one-dimensional reduction of the overall physiological state that reflects the system’s capacity to maintain stability. Dynamic changes of *x*, accompanied by the resulting alteration of the internal parameters of S and W waves, can be initiated by homeostatic and other factors, such as the circadian clock or environmental perturbations.

The probabilistic nature of both S and W waves allows us to employ the well-developed mathematical framework of quantum mechanics to quantitatively describe the transitions between these two states.

Our prior investigations yielded successful results through this approach, providing precise quantitative descriptions of normal human sleep architecture by solving the wave equation for the *x*-variable^16^. For the examination of the 24-hour dynamics of S-W transitions in this study, we adopted a simpler semiclassical approach^28,29^. This method relies on the classical-like motion of the center of a wave packet corresponding to the S or W wave. Within this approximation, the regulating parameter x is treated as a predetermined function of time, denoted as x=x(t), and the evolution and transitions take place due to the dynamic adjustments in *x*.

#### Semi-classical evolution of the two-state system

Within the framework of the wave model, the time-evolution of the two-state system can be described by the superposition of sleep and wake waves:

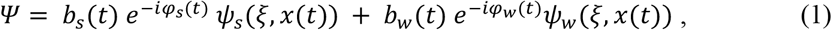

where *b*_*S*_(*t*) and *b*_*W*_(*t*) are amplitudes of the S-and W-waves and

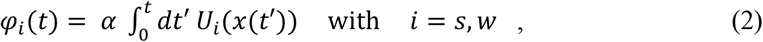

where *φ* is the phase of S or W wave accumulated since wakeup time at *t=0*. The constant *α* is introduced for the transformation of the unitless time *t* to hours. The potential energies *U*_*S*_(*x*(*t*)) and *U*_*W*_(*x*(*t*)) modify the propagation of S and W waves, respectively, or the motion of their wavepacket, and reflect the degree of state instability.

The numerical parameters of the *U*_*S*_ and *U*_*W*_ potentials were obtained from the detailed analysis of sleep architecture in data collected by our group (see above) and data reported by Barbato and Wehr^30^. The results of the analysis are described in detail in^16^. The established potential curves *U*_*S*_ and *U*_*W*_ provided the basis for the present study, allowing us to determine how the energy of state instability changes depending on the 24-h variation in the regulating parameter x(t).

The time-dependent amplitudes *b*_*S*_(*t*) and *b*_*W*_(*t*) of S and W waves provide the time-dependent probabilities of being in S or W state:

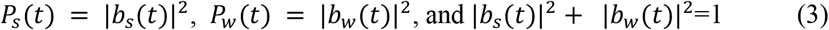

The equations governing the two-state dynamics with time-dependent parameters have been the subject of thorough exploration in quantum physics. Examples include analysis of populations of excited states in atomic collisions^28^, the experimental examination of the state transitions in ultracold gases^31^ and the investigation of atomic state populations under the influence of time-dependent electric fields^32^. The equations corresponding to the amplitudes of S and W states were derived from the wave equation governing the total wave function *Ψ*. This was accomplished through the conventional process of projecting the time-dependent wave equation onto the basis functions *ψ*_*S*_ and *ψ*_*W*_ ^29^:

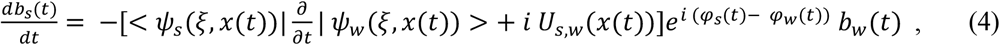

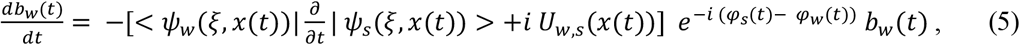

where 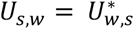 is the matrix element describing the potential of interaction between S and W states. In our model, the value of *U*_*S,W*_ includes an explicit dependence on *x* only, therefore the time variation of *x* = *x*(*t*) changes S-W interaction. The value of *U*_*S,W*_(x) peaks in the vicinity of the crossing point *x* = *x*_*c*_ and we use the Gaussian shape of this peak to simplify the calculations: *U*_*S,W*_(*x*) = *γ exp*[− (*x* − *x*_*c*_)^2^/2 *β*^2^], where parameters *γ* and *β* reflect the maximal value of the interaction matrix element and the width of the interaction area, respectively. Under real conditions, *U*_*S,W*_ may also include an explicit dependence on time *t* due to circadian and environmental factors, *U*_*S,W*_ = *U*_*S,W*_(*x, t*).

The knowledge of the time-dependence of the regulating parameter *x=x(t)* allows us to rewrite Eqs. 4 and 5 in the form convenient for numerical solutions:

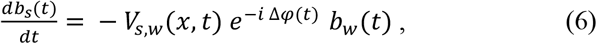

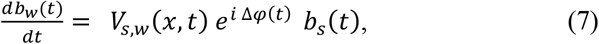

where *ν*(*t*) = *dx*(*t*)/*dt* is the velocity associated with the transformation of the regulating parameter during the 24-hour S-W cycle. The difference of S and W phases Δ*φ*(*t*) = *φ*_*W*_(*t*) − *φ*_*S*_(*t*) in Eqs. 6 and 7 depends on the energy separation Δ*U*(*x*) = *U*_*W*_(*x*) − *U*_*S*_(*x*) and the velocity of *x, ν*(*t*):

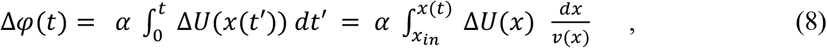

where Δ*φ*(*t*) must be calculated for the known trajectory *x*=*x*(*t*) or for the velocity *ν* = *ν*(*x*), which is defined as the function of *x*. In Eqs. 6 and 7, the matrix element *V*_*S,W*_(*x, t*) includes the contributions of two possible mechanisms of interaction between S and W waves:

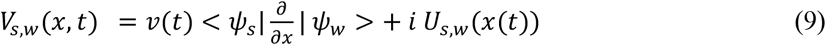

#### Analysis of Sleep-Wake interaction and Sleep Propensity

Equations 6 and 7 describe the dynamics of the amplitudes *b*_*S*_(*t*) and *b*_*W*_(*t*) corresponding to the S and W waves. As a result, they offer insight into the time-dependent probability of remaining awake or transitioning into sleep. These dynamics of *b*_*S*_(*t*) and *b*_*W*_(*t*) also reflect the effectiveness of the S-W interaction across various regions of the regulating parameter *x*. The mathematical characteristics of Eqs. 6 and 7 allow to predict the region where efficient transitions between S and W can take place. The rate of S ↔ W transitions can be notably elevated within regions characterized by slow alterations of the phase difference Δφ(t) or within domains exhibiting elevated values of *V*_*S,W*_(*x, t*). In alignment with the principles of the theory of non-adiabatic transitions^26,27^ these criteria can be succinctly expressed through Massey’s parameter Z:

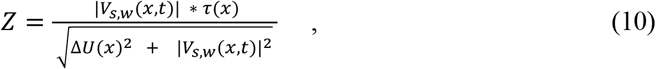

where *τ*(*x*) is the characteristic time of the S-W interaction at certain *x*. All values in Eq. 10 are expressed in dimensionless units. The typical dependence of *Z* on the regulating parameter *x* is shown in Fig. 4b, with *τ* assumed constant for simplicity. Maximal interaction strength corresponds to the region of the crossing of the *U*_*W*_(*x*) and *U*_*W*_(*x*) potential curves, where |Δ*U*(*x*)| ≪ |*V*_*S,W*_(*x, t*)|, as illustrated in Fig. 4b.

The primary objective of this study was to formulate accurate theoretical predictions for the daily variations in sleep propensity, which can be understood as the rate of transitions from Wake to Sleep states at different times *t* throughout the 24-hour cycle. This transition rate was quantified by calculating the probability *P*_*W,S*_(*t*, Δ*t*) of transitioning from wake into sleep within a fixed time interval Δt. To prevent significant alterations in homeostatic parameters during this interval, Δt was selected to be sufficiently small. For our investigation, the value of Δt was set at 0.1 hours or 6 minutes.

We computed *P*_*W,S*_(*t*, Δ*t*) for the time span 0 ≤ *t* ≤ 24hours, where t=0 corresponds to the habitual morning awakening time. The temporal trajectory of the regulating parameter *x=x(t)* was determined using a series of reference points established in our previous study. These model points were inferred from an extensive quantitative analysis of human sleep architecture within specific experimental groups with regular and extended sleep^16^ (Fig. 4a).

#### Probability of state transitions in the absence of sleep

First, we carried out calculations to determine the probability of state transitions over a 24-hour interval in the absence of sleep. Utilizing the established time-dependent value of the regulating parameter *x(t)*, we were able to derive the time-evolution of the instability energy ε(t) as follows: *ε(t) = U*_*w*_*(x(t))*. The outcomes of these calculations are depicted in Fig. 5a, illustrating the variations of *ε(t)* attributed to the variable rate of progression of the regulating parameter *x*. The initial conditions required for the determination of the 24-hour sleep propensity *P*_*W,S*_(*t*, Δ*t*) are: *b*_*W*_(*t*) = 1; *b*_*S*_(*t*) = 0 and *P*_*W,S*_(*t*, Δ*t*) = |*b*_*S*_(*t* + Δ*t*)|^2^ for any *t* ≤ 24. The calculated rate of transition in *P*_*W,S*_(*t*, Δ*t*) is depicted in Fig. 5b

#### Probability of state transitions in the presence of overnight sleep

The analysis of the probability of state transitions within a 24-hour timeframe, which encompassed an 8-hour overnight sleep, consisted of two distinct stages (Fig. 5c). The initial step involved the computation of 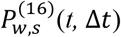 throughout the 16-hour period of wakefulness, where 0 ≤ *t* ≤ 16. The superscript index “16” indicates the duration of the continues wakefulness period.This calculation employed the same *x(t)* trajectory and parameters as detailed earlier, in the absence of sleep, with 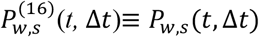 (Fig. 5).

During the 8-hour sleep interval, encompassing the time period 16 ≤ *t* ≤ 24, the evaluation of sleep propensity followed different initial conditions, with *b*_*S*_(*t*) = 1; *b*_*W*_(*t*) = 0. These conditions corresponded to the transition speed from S to W state, resulting in the calculation of 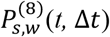. The straightforward relationship between the probability of staying asleep and the probability of waking up during sleep, 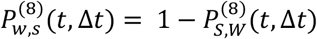, allows for the theoretical evaluation of the sleep propensity during 8-hour sleep.

The quantitative description of sleep architecture through the wave model^16^ and the determination of the durations of consecutive NREM and REM episodes facilitated the computation of 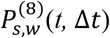 using a theoretical formula for the probability of remaining asleep after undergoing “*n*” sleep cycles:

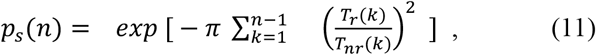

where *T*_*r*_(*k*) and *T*_*nr*_(*k*) are the durations of REM and NREM episodes in the k-th sleep cycle. The formula in Eq. 11 has been derived with the assumption of Landau -Zener’s type of interaction^32^ between S and W states in the vicinity of the *x* = *x*_*c*_ region where the two potential curves cross (Fig. 2, 4a). Since the model accurately predicts *T*_*r*_(*k*) and *T*_*nr*_(*k*)^16^, in the Eq. 11 we used either experimental or theoretical data to calculate *p*_*S*_(*n*), with similar results. The speed of W-S transition for the two time intervals, 16 and 8 hours, was matched at t=16 hours (Fig. 5c, black line).

In experimental settings, alterations in wake propensity *P*_*S,W*_ during the sleep period can be approximated by directly measuring the percentage of time spent awake^17^. Analogously to the previously discussed method, this parameter can be transformed into an estimation of sleep propensity using the straightforward relationship *P*_*W,S*_ = 1 − *P*_*S,W*_. The outcomes of employing this approach are depicted in Figure 4c (red dashed line) and exhibit similarities with those yielded by Eq. 11.

### Data availability statement

The datasets generated by the co-author (IVZ and AAP) and analyzed during the current study are available from the corresponding authors on reasonable request.

The mathematical algorithms of the wave model of sleep dynamics in “Wolfram Mathematica” format are available from the corresponding author (VK) on reasonable request.

## Institutional Review Board Statement

The experimental studies were conducted in accordance with the Declaration of Helsinki, and approved by the Institutional Review Board of Boston University School of Medicine (protocol code H-33035 and date of approval is November 15, 2015) and the Ethics Committee of the Siberian Branch of the Russian Academy of Medical Sciences (2010).

## Author contributions

V.K., I.V.Z. and A.A.P. designed research; I.V.Z and A.A.P. performed sleep research; V.K. and M.R. conducted mathematical modeling; V.K. and I.V.Z. wrote the paper.

